# Evaluation of Enterobacterales carrying *Acinetobacter*-associated *bla*_OXA_ genes—United States, 2017–2022

**DOI:** 10.1101/2025.06.05.655513

**Authors:** Giulia Orazi, Alyssa G. Kent, Nadine Wilmott, Desbelet Berhe, Cynthia Longo, Porscha Bumpus-White, Debra Rutledge, Gregory Hovan, Ilsa M. Villegas Correa, Inelia Otero Pagán, Penny Coe, Michelle Therrien, Amelia Bhatnagar, Susannah L. McKay, Allison C Brown, Alison Laufer Halpin, Sarah Sabour

## Abstract

Through their ability to hydrolyze carbapenems, Ambler class D beta-lactamases endanger patients by limiting the clinical efficacy of beta-lactam antimicrobials. Further, plasmid-mediated transmission can increase mobility of carbapenemase genes between bacteria and facilitate their spread between patients. In the United States and elsewhere, the plasmid-mediated Ambler class D carbapenemase genes *bla*_OXA-23-like_, *bla*_OXA-24/40-like_, and *bla*_OXA-58-like_ are commonly associated with *Acinetobacter* species and have rarely been reported outside of this genus. However, multiple recent international reports indicate detection of Enterobacterales isolates carrying *Acinetobacter*-associated class D carbapenemase genes. This evaluation aimed to provide insight into whether Enterobacterales harboring these class D genes may be circulating undetected in the United States, thereby signaling a need for adapting testing strategies to prioritize the detection of these potentially emerging public health threats. We analyzed whole-genome sequencing data generated through multiple Centers for Disease Control and Prevention (CDC) activities, including testing conducted across the Antimicrobial Resistance Laboratory Network, to determine whether any Enterobacterales isolates sequenced from 2017–2022 harbored *Acinetobacter*-associated class D carbapenemase genes. Among ∼10,000 predominantly carbapenem-resistant Enterobacterales isolates, we identified only a single Enterobacterales isolate harboring an *Acinetobacter*-associated class D gene – a *Klebsiella pneumoniae* isolate harboring a *bla*_OXA-23-like_ gene. Our findings suggest that *bla*_OXA-23-like_, *bla*_OXA-24/40-like_, and *bla*_OXA-58-like_ genes are rare among Enterobacterales isolates sequenced through CDC public health activities in the United States and do not warrant changes to current testing priorities at this time.

## Introduction

The continued global spread of carbapenem resistance threatens our collective ability to treat and prevent severe infections caused by Gram-negative bacterial pathogens. In particular, carbapenem-resistant *Acinetobacter* and Enterobacterales are two of the five public health antibiotic resistance threats determined by the Centers for Disease Control and Prevention (CDC) as requiring “urgent and aggressive action” (1). These pathogens can cause serious infections that are often associated with poor patient outcomes, including treatment failure and death (2–5).

It is well established that a variety of carbapenemases contribute to carbapenem resistance in Enterobacterales (5, 6). In the United States, the most common carbapenemase-encoding genes detected among healthcare-associated carbapenem-resistant Enterobacterales (CRE) are: *bla*_KPC_, *bla*_NDM_, *bla*_OXA-48-like_, *bla*_VIM_, and *bla*_IMP_ (7, 8). The Antimicrobial Resistance Laboratory Network (AR Lab Network) is a network of United States public health laboratories (PHLs) that performs molecular testing to detect these five targeted carbapenemase genes in CRE. From 2017–2023, over one-third of CRE isolates tested by the AR Lab Network (35.4% [n=42,072/118,748]) harbored at least one of these carbapenemase genes (8). The most common gene detected in CRE isolates was *bla*_KPC_ (26.0%), followed by *bla*_NDM_ (7.7%), while the three other genes were less commonly detected (<2% each) (8).

In contrast, the most common mechanisms conferring carbapenem resistance in *A. baumannii* globally are the acquired Ambler class D beta-lactamases OXA-23, OXA-24/40, and OXA-58 (9), which are among the oxacillinases or OXA-type carbapenemases. Despite exhibiting weak carbapenem-hydrolyzing activities compared to other carbapenemases, these enzymes can render treatment ineffective (10, 11). A recent United States evaluation of AR Lab Network-generated data reported that 83% (n=3,351/4,041) of carbapenem-resistant *A. baumannii* (CRAB) isolates originating from 45 states between 2017–2020 tested positive for at least one of these acquired OXA-type carbapenemase genes (12).

Although historically *bla*_OXA-23-like_, *bla*_OXA-24/40-like_, and *bla*_OXA-58-like_ genes have been limited to *Acinetobacter* species, carbapenemase-encoding genes can be horizontally acquired and disseminated between organisms. For example, plasmids can mediate the exchange of carbapenemase genes between different bacterial strains, species, and genera (13) and, as a result, facilitate the spread of resistance to this important class of antimicrobials between patients and across healthcare facilities (14). Several international reports indicate these *Acinetobacter*-associated class D genes can also be found in members of the order Enterobacterales: *Enterobacter cloacae* (15), *Klebsiella pneumoniae* (15–17), *Escherichia coli* (15, 16, 18), and most commonly, *Proteus mirabilis* (19–25).

To our knowledge, only one previous report has identified any Enterobacterales isolates in the United States carrying these genes – 10 *K. pneumoniae* isolates belonging to various sequence types that harbored either *bla*_OXA-23_ (n=5) or *bla*_OXA-24/40_ (n=5) (17). These genes were detected through a retrospective, population-based analysis of *K. pneumoniae* isolated from patients treated at a single hospital system in Houston, Texas, between 2011 and 2015. Because *bla*_OXA-23-like_, *bla*_OXA-24/40-like_, and *bla*_OXA-58-like_ genes have been predominantly associated with *Acinetobacter* species, AR Lab Network testing for these genes is performed exclusively on *Acinetobacter* spp. isolates and CDC activities have yet to assess the occurrence of these genes among Enterobacterales isolates collected across the United States. To our knowledge, no such analyses have been previously conducted in the United States on a large scale.

Investigating combinations of carbapenemase genes and organisms that are not currently targeted is essential for uncovering hidden reservoirs of carbapenemase genes and informing testing priorities and strategies for rapidly detecting emerging public health threats. We performed an evaluation using a large, national collection of contemporary isolates with available whole-genome sequencing (WGS) data to investigate whether Enterobacterales isolates sequenced through CDC multiple activities harbor *Acinetobacter*-associated class D carbapenemase genes as has been reported internationally.

## Materials and Methods

### Retrospective analysis of WGS data from Enterobacterales isolates collected through multiple CDC activities

A dataset of short-read WGS data for ∼10,000 Enterobacterales isolates collected through CDC’s healthcare-associated infections surveillance activities (Emerging Infections Program and Division of Healthcare Quality Promotion Sentinel Surveillance system), outbreak investigations, reference testing, and AR Lab Network activities from 2017–2022 was reviewed retrospectively for the presence of *Acinetobacter*-associated class D genes (**Fig. S1**).

The current state of our data infrastructure and challenges in linking data from different sources makes it infeasible to determine what proportion of these sequences are duplicates due to resequencing, and thus provide precise denominators for isolates sequenced through each activity. The WGS dataset included Illumina sequencing data available for isolates of three more common Enterobacterales genera (*Enterobacter, Escherichia*, and *Klebsiella*) and less common Enterobacterales genera (*Citrobacter, Hafnia, Morganella, Pantoea, Proteus, Providencia, Raoultella*, and *Serratia*) (7, 26).

Out of the ∼10,000 total Enterobacterales isolate sequences in the WGS dataset, ∼4,700 were generated through the AR Lab Network, which targets carbapenem-resistant isolates (**Fig. S1**). This WGS dataset likely includes both carbapenem-resistant and carbapenem-susceptible isolates, and not all isolates sequenced as part of the CDC activities above underwent antimicrobial susceptibility testing (AST). As noted above, a proportion of isolate sequences included in this dataset may be duplicate sequences due to resequencing of the same isolate; challenges in data infrastructure and data linking hamper deduplication efforts.

Sequences in the WGS dataset were analyzed using CDC’s in-house bioinformatics pipeline (QuAISAR-H, https://github.com/DHQP/QuAISAR_singularity). QuAISAR-H was used to perform quality control of Illumina sequencing data, genome assembly, and annotate antimicrobial resistance genes using a custom in-house database. WGS data were searched for the presence of all previously identified alleles of *bla*_OXA-23-like,_ *bla*_OXA-24/40-like_, and *bla*_OXA-58-like_ genes (27) using the following thresholds: 98% amino acid identity and 90% coverage over the length of the respective allele.

PlasFlow v1.1 (28) and Unicycler v0.4.4 (29) were used to identify plasmids. Plasmid typing was performed using PlasmidFinder v1.3 (30). BLASTn (BLAST+ v2.13.0) (31) was used to identify the best match of the *bla*_OXA-23-like_ gene-containing contig to the NCBI nucleotide database (32) based on percent coverage and identity. Identification of any of the above genes through this retrospective WGS analysis was then confirmed at CDC by conducting real-time-PCR (RT-PCR) on the stored bacterial isolate as previously described (7, 12). BRIG (33) was used to visualize and compare plasmids.

### Characterization of Enterobacterales isolates collected through the AR Lab Network

As part of the AR Lab Network, PHLs in all 50 states, five large cities, and Puerto Rico characterize clinical isolates of CRE submitted by a network of clinical laboratories in their jurisdiction (7). As previously described, PHLs perform organism identification, phenotypic characterization (AST and carbapenemase production testing), and molecular testing of isolates initially detected as carbapenem-resistant by submitting clinical laboratories (7). All carbapenemase-production-positive CRE isolates (n=94,186 during 2017–2022) underwent RT-PCR testing for detection of five targeted carbapenemase genes (*bla*_KPC_, *bla*_NDM_, *bla*_OXA-48-like_, *bla*_IMP_, and *bla*_VIM_) (7, 12) (**Fig. S1**). Unlike CRAB isolates collected through the AR Lab Network, which also undergo additional RT-PCR testing for *Acinetobacter*-associated class D carbapenemase genes (*bla*_OXA-23-like_, *bla*_OXA-24-like_, and *bla*_OXA-58-like_) (12), CRE isolates are not tested for the presence of these genes.

Either AR Lab Network PHLs or CDC perform WGS on a subset of CRE isolates, including isolates that are either carbapenemase-production-positive or RT-PCR-positive for a targeted carbapenemase gene, isolates resistant to all tested antimicrobials, and isolates identified as part of an outbreak. CDC performs supplemental AST on CRE isolates with rare or novel phenotypic or genotypic profiles, including the *bla*_OXA-23-like_-harboring *K. pneumoniae* isolate identified through this evaluation, using reference broth microdilution as previously described (34). AST results are interpreted using Clinical and Laboratory Standards Institute (CLSI) M100 breakpoints (35); for antimicrobials with no CLSI breakpoints available, U.S. Food and Drug Administration breakpoints are used (36).

### Analysis of *bla*_OXA-48-like_ gene frequency among CRE isolates tested in the AR Lab Network

Enterobacterales are more commonly found to harbor *bla*_OXA-48-like_ genes (7, 8, 26, 37–41). To contextualize the findings of this evaluation on the frequency of *Acinetobacter*-associated class D genes in CRE, RT-PCR data generated by PHLs during routine testing were used to determine the frequency of detection of *bla*_OXA-48-like_ genes among CRE isolates tested in the AR Lab Network during 2017–2022 (n=94,186). Frequencies were calculated in aggregate and by organism groupings as follows: 1) all Enterobacterales, 2) *E. coli*, 3) *Klebsiella* spp., 4) *Enterobacter* spp., and 5) other, less common Enterobacterales genera, as defined above. Only isolates identified at the genus level were included in this analysis.

### Data availability

Whole-genome sequencing data generated by CDC and other public health laboratories are publicly available at the BioProject [PRJNA288601](https://www.ncbi.nlm.nih.gov/bioproject/288601) (Gram-negative bacteria), which is within the umbrella project [PRJNA531911](https://www.ncbi.nlm.nih.gov/bioproject/531911) (all healthcare-associated infections). Whole-genome sequencing data of the *Klebsiella pneumoniae* isolate were generated in 2017 and are publicly available as follows: this Whole Genome Shotgun project has been deposited at GenBank under the accession JBCPYB000000000. The version described in this paper is version JBCPYB010000000. Data resulting from bioinformatic analyses of the *K. pneumoniae* isolate are included in this article and supplemental files. Molecular detection data generated through the Antimicrobial Resistance Laboratory Network and analyzed or referenced in this article are publicly available on CDC’s [Antibiotic Resistance & Patient Safety Portal](https://arpsp.cdc.gov/profile/antibiotic-resistance?tab=ar-lab-network).

## Results

Through a retrospective analysis of WGS data on ∼10,000 Enterobacterales isolates, we identified a single isolate harboring an *Acinetobacter*-associated class D carbapenemase gene: a *K. pneumoniae* isolate (ST 896) harboring a *bla*_OXA-23-like_ gene. The isolate was one of ∼4,700 Enterobacterales isolates collected and sequenced through the AR Lab Network from 2017– 2022 and was originally cultured from a blood specimen in 2017.

AST results showed the isolate was resistant to 17 of 24 antimicrobials tested, corresponding to seven classes, including carbapenems and cephalosporins (**Table 1**). The isolate was positive for phenotypic carbapenemase production but RT-PCR-negative for all five targeted carbapenemase genes, suggesting the possible presence of a non-targeted carbapenemase gene or variant.

**Table 1.** Antimicrobial susceptibility testing profile of the *bla*_OXA-23-like_-harboring *Klebsiella pneumoniae* isolate.

| Antimicrobial class | Antimicrobial | MIC (µg/ml) | Interpretation |
| --- | --- | --- | --- |
| Carbapenem | Ertapenem | >8 | R |
|  | Meropenem | >8 | R |
|  | Imipenem | 32 | R |
|  | Doripenem | >8 | R |
| Cephalosporin | Cefotaxime | >64 | R |
|  | Ceftazidime | >128 | R |
|  | Ceftriaxone | >32 | R |
|  | Cefepime | >32 | R |
|  | Cefoxitin | >16 | R |
|  | Cefazolin | >8 | R |
| Monobactam | Aztreonam | >64 | R |
| Beta-lactam combination | Piperacillin-tazobactam | >128/4 | R |
| Penicillin | Ampicillin | >32 | R |
| Tetracycline | Tetracycline | >32 | R |
|  | Tigecycline | ≤0.5 | S |
| Antifolate combination | Trimethoprim-sulfamethoxazole | >8/152 | R |
| Fluoroquinolone | Ciprofloxacin | 2 | R |
|  | Levofloxacin | 2 | R |
| Aminoglycoside | Amikacin | 2 | S |
|  | Gentamicin | 0.5 | S |
|  | Tobramycin | ≤0.5 | S |
| Amphenicol | Chloramphenicol | 8 | S |
|  | Colistin | 0.5 | I |

Antimicrobial susceptibility testing was performed on a *K. pneumoniae* isolate that was collected through the Antimicrobial Resistance Laboratory Network in 2017. Antimicrobial susceptibility testing was performed at CDC using reference broth microdilution and results were interpreted according to CLSI document M100 34 ed. (32), with the exception of tigecycline, for which criteria provided by the U.S. Food and Drug Administration were applied. Abbreviations: MIC, minimum inhibitory concentration; CLSI, Clinical and Laboratory Standards Institute; R, Resistant; I, Intermediate; S, Susceptible.

Analysis of WGS data identified 19 AR genes in this isolate (**Table S1**), including four beta-lactamase genes: *bla*_SHV-12_, *bla*_TEM-212_, *bla*_LAP-2_, and *bla*_OXA-1095_. *bla*_OXA-1095_ is a novel variant that differs by a single amino acid substitution from *bla*_OXA-23_ (GenBank accession: OM933714.1; RefSeq accession: NG_079932.1) (27). WGS data from this evaluation suggest that *bla*_OXA-1095_ is located on the *Acinetobacter* resistance island AbaR4, specifically within the transposon Tn2006 (**Fig. 1**). The rest of the 49-kb contig (**Fig. 1**) matched a previously identified, non-typeable *Klebsiella* plasmid, pKPN-704 (42) carrying multiple genes encoding conjugation machinery (99% nucleotide identity; 97% coverage). The other beta-lactamase genes identified in the isolate are not located on the same plasmid as *bla*_OXA-1095_.

**Figure 1.**
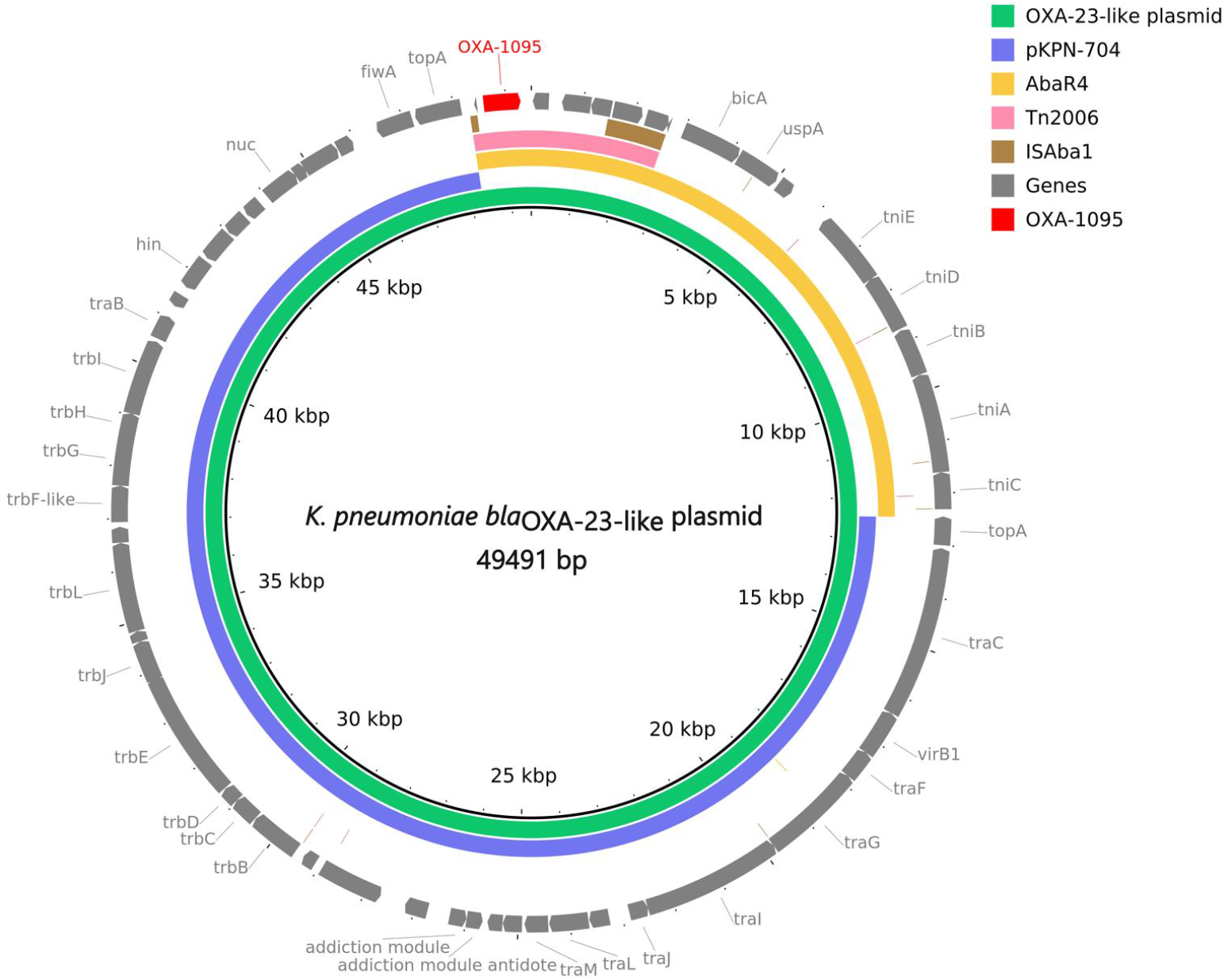
Short-read whole-genome sequencing data of the *bla*_OXA-23-like_-harboring *Klebsiella pneumoniae* isolate suggest that *bla*_OXA-1095_ is located on a nested mobile genetic element composed of the *Acinetobacter* resistance island AbaR4 and a non-typeable *K. pneumoniae* plasmid. Among ∼10,000 Enterobacterales isolates sequenced through CDC activities from 2017–2022, a single isolate was found to harbor an *Acinetobacter*-associated class D gene. Whole-genome sequencing data were generated for this *K. pneumoniae* isolate that was collected through the Antimicrobial Resistance Laboratory Network in 2017. The innermost, green ring represents the sequence of the *bla*_OXA-1095_-harboring plasmid (GenBank accession number: JBCPYB000000000). The surrounding colored rings indicate the most similar sequences in the NCBI nucleotide database, including the plasmid pKPN-704 (GenBank accession number: CP014764.1); all hits shown have >99% nucleotide sequence identity. In the outermost ring, *bla*_OXA-1095_ is shown in red and other annotated ORFs are shown in grey.

To contextualize the rarity of *Acinetobacter*-associated class D genes in Enterobacterales, we determined the frequency of another Ambler class D gene, *bla*_OXA-48-like_, among CRE isolates tested in the AR Lab Network during the same time period (2017–2022) using RT-PCR data. We found that 1.52% of CRE isolates (n=1,436/94,186) harbored *bla*_OXA-48-like_ genes. The frequency of *bla*_OXA-48-like_ detection in CRE varied by genus: *E. coli* and *Klebsiella* spp. isolates exhibited similar frequencies of 3.36% (n=482/14,344) and 2.3% (n=909/39,470), respectively, whereas only 0.04% (n=11/28,049) of *Enterobacter* spp. isolates were found to carry these genes. Collectively, less common Enterobacterales genera (7, 26) harbored these genes at a frequency of 0.28% (n=34/12,323).

## Discussion

Recent evidence worldwide suggests that certain class D carbapenemase genes (i.e., *bla*_OXA-23-like_, *bla*_OXA-24/40-like_, and *bla*_OXA-58-like_ genes) have spread from *Acinetobacter* to Enterobacterales. However, due to the absence of routine molecular testing for the presence of these genes in Enterobacterales, it is unclear how common these genes are among Enterobacterales strains circulating in the United States. Through our retrospective evaluation of available data collected on a contemporary collection of thousands of Enterobacterales isolates sequenced across a variety of nationwide CDC public health activities, we identified a single *K. pneumoniae* isolate carrying a *bla*_OXA-23-like_ gene. To our knowledge, the only other report of these genes in Enterobacterales isolated in the United States describes their detection at a single hospital system and predates the establishment of the AR Lab Network (17). Together, these findings suggest these *Acinetobacter*-associated class D genes are rare among CRE in the United States.

The *K. pneumoniae* isolate we identified through this evaluation harbored a novel *bla*_OXA-23-like_ variant that was designated as *bla*_OXA-1095_. AST showed the isolate was resistant to most of the antimicrobials tested, including carbapenems, traditional beta-lactams, and fluoroquinolones, which is consistent with AST profiles of *K. pneumoniae* and *E. coli* isolates previously reported to carry *bla*_OXA-23_ (16, 18, 43, 44). Because multiple beta-lactamase genes were detected in the isolate from this evaluation and those from previous reports, the precise contributions of *bla*_OXA-1095_ and other *bla*_OXA-23-like_ genes to the AST phenotypes of these organisms are unknown.

Our data suggest that *bla*_OXA-1095_ is located on a composite mobile genetic element consisting of a *K. pneumoniae* plasmid and the *Acinetobacter* resistance island AbaR4. A similar genetic context (i.e., plasmid-localized AbaR4) has been observed for *bla*_OXA-23_ in strains of *A. baumannii* (9, 45) and in a *P. mirabilis* isolate from Singapore (46). Given that AbaR4 is thought to rely on plasmids for interspecies transfer (46, 47), and *Acinetobacter* plasmids have been identified in Enterobacterales (48, 49), plasmids may have enabled the transfer of *bla*_OXA-23-like_ genes between these organisms.

Plasmids may also be facilitating the spread of class D carbapenemase genes from Enterobacterales to *Acinetobacter* (50). Recently, the AR Lab Network detected CRAB isolates harboring *bla*_OXA-48-like_ genes, a group of class D genes associated with Enterobacterales (7, 8, 26, 37–41) that to our knowledge had not been observed in CRAB previously; five *bla*_OXA-48-like_-positive CRAB isolates were detected in 2021 and ten in 2022 (8). The dissemination of carbapenem resistance mechanisms to additional bacterial taxa threatens to compromise the efficacy of clinically important antimicrobials against multiple human pathogens and challenges our ability to prevent the spread of these genes within healthcare settings and beyond.

The findings from this evaluation suggest that *Acinetobacter*-associated class D carbapenemase genes, which are not targeted for molecular detection (by RT-PCR) in Enterobacterales by the AR Lab Network or other CDC activities, are extremely rare among Enterobacterales isolates collected across the United States through current public health activities. In contrast, *bla*_OXA-48-like_ genes are more common among Enterobacterales isolates in the United States and worldwide (7, 8, 26, 37–41), and are routinely tested in these organisms by AR Lab Network PHLs. Among our large contemporary collection of ∼90,000 CRE isolates tested through the AR Lab Network (8), *bla*_OXA-48-like_ genes were detected at a frequency of 1.52% – two orders of magnitude higher than *Acinetobacter*-associated class D genes (detected in one out of ∼10,000 isolates).

This evaluation had two important limitations. First, this collection of isolates is biased towards carbapenem-resistant phenotypes, and second, AST was not performed on all isolates represented in the WGS dataset. Among ∼10,000 isolates in this collection, ∼4,700 isolates were collected through the AR Lab Network, which selects isolates on the basis of carbapenem resistance. Specifically, isolate testing through the AR Lab Network is initiated when carbapenem resistance is detected by a submitting clinical laboratory. A large proportion of isolates collected through surveillance activities as part of the Emerging Infections Program were also targeted based on carbapenem-resistant case definitions, and outbreak investigations are frequently triggered by detection of carbapenemase production.

When testing priorities target carbapenem-resistant phenotypes, class D carbapenemase genes may escape detection, as some Enterobacterales isolates carrying these genes have been reported to be susceptible to carbapenems. For example, international reports identified several *P. mirabilis* isolates harboring *bla*_OXA-23_ that tested susceptible or borderline susceptible to ertapenem and meropenem (19, 20, 23, 24). Additionally, in the United States, a portion of *bla*_OXA-48-like_-positive CRE isolates tested in the AR Lab Network from 2017–2022 were susceptible to doripenem (43%), meropenem (39%), or imipenem (31%), compared to only 3% of isolates susceptible to ertapenem (8). Therefore, the focus of AR Lab Network and other CDC activities on carbapenem resistance may underestimate the number of Enterobacterales isolates harboring these genes that are circulating in the United States.

An additional limitation is that the repository of isolates collected and sequenced through CDC activities may not be representative of all clinical Enterobacterales isolates across the United States for multiple reasons. There may be uneven representation across geographic regions due to jurisdictional differences in regulations and/or isolate submissions to the AR Lab Network and participation of a limited number of sites in surveillance activities (e.g., the Emerging Infections Program). The AR Lab Network is not a surveillance system, but a laboratory response network created to rapidly detect and contain antimicrobial resistance threats; the exploration of data for surveillance purposes is a secondary goal of the Network.

Finally, despite deduplication of data where possible, some isolates may have been submitted and/or sequenced more than once for different activities, including reference testing and outbreak investigations. Due to the current state of our data infrastructure and challenges in linking data from different sources, it is not feasible to determine what proportion of these sequences are duplicates due to resequencing, and thus provide precise denominators for isolates sequenced through each activity. We have included a conservative denominator estimate of ∼10,000 to avoid providing a more exact number that may be misleading.

## Conclusions

Through retrospective analysis of WGS data, this evaluation indicated that *Acinetobacter*-associated class D carbapenemase genes were extremely rare among Enterobacterales isolates sequenced through multiple CDC public health activities (detected in only one out of ∼10,000 isolates). Our findings suggest that these *Acinetobacter*-associated carbapenemase genes are uncommon among CRE isolates in the United States currently and do not warrant additional resources for routine detection at this time. However, Enterobacterales species – including carbapenem-susceptible populations that are not currently monitored by public health activities – could represent potential reservoirs for the spread of these carbapenemase genes. To uncover hidden reservoirs of carbapenemase genes and understand their frequency and clinical implications, it is essential to evaluate gene-organism combinations beyond those that are routinely targeted for detection. Performing these types of investigations is critical for guiding detection priorities and assay development for the prevention and containment of emerging antimicrobial resistance threats in healthcare.

## List of abbreviations

AR Lab Network: Antimicrobial Resistance Laboratory Network
CDC: Centers for Disease Control and Prevention
CRAB: carbapenem-resistant *Acinetobacter baumannii*
CRE: carbapenem-resistant Enterobacterales
PHLs: public health laboratories

## Ethics approval

This activity was reviewed and approved by the CDC as exempt human subjects research (#7218, “Use of Residual Samples for Public-Health Research and Development to Prevent Healthcare-Associated Infections” (45 CFR 46.104(d)(4ii))).

## Funding

This work is supported by Cooperative Agreement Number NU60OE000104 (CFDA #93.322), funded by the Centers for Disease Control and Prevention of the US Department of Health and Human Services. This project was 100% funded with federal funds from a federal program of $120,402,978.

### Disclaimer

The findings and conclusions in this report are those of the authors and do not represent the official position of the Centers for Disease Control and Prevention, the US Department of Health and Human Services, or the Association of Public Health Laboratories.

## Author contributions

All authors contributed to at least one of the following: conception and design (SS, ACB, AB, GO), acquisition of bacterial isolates and laboratory testing (NW, CL, PBW, DR, GR, IMVC, IOP, PC, MT), analysis and interpretation of laboratory data sets (DB, GO), bioinformatic analysis and interpretation of sequencing data and data sets (AGK), preparation of figures and tables (AGK, GO), drafting the manuscript (GO), and substantive review/revision of the manuscript (SS, ACB, SLM, ALH, AB, GO). All authors read, reviewed, and approved the final manuscript.

## References

1. CDC. Antibiotic resistance threats in the United States, 2019. https://www.cdc.gov/drugresistance/pdf/threats-report/2019-ar-threats-report-508.pdf.

2. Schwaber MJ, Klarfeld-Lidji S, Navon-Venezia S, Schwartz D, Leavitt A, Carmeli Y. 2008. Predictors of carbapenem-resistant Klebsiella pneumoniae acquisition among hospitalized adults and effect of acquisition on mortality. Antimicrobial Agents and Chemotherapy 52:1028–1033.

3. Akova M, Daikos GL, Tzouvelekis L, Carmeli Y. 2012. Interventional strategies and current clinical experience with carbapenemase-producing Gram-negative bacteria. Clinical Microbiology and Infection 18:439–448.

4. Weiner LM, Webb AK, Limbago B, Dudeck MA, Patel J, Kallen AJ, Edwards JR, Sievert DM. 2016. Antimicrobial-resistant pathogens associated with healthcare-associated infections: summary of data reported to the National Healthcare Safety Network at the Centers for Disease Control and Prevention, 2011–2014. Infection Control & Hospital Epidemiology 37:1288–1301.

5. Jean S-S, Harnod D, Hsueh P-R. 2022. Global threat of carbapenem-resistant Gram-negative bacteria. Frontiers in Cellular and Infection Microbiology 12.

6. Ma J, Song X, Li M, Yu Z, Cheng W, Yu Z, Zhang W, Zhang Y, Shen A, Sun H, Li L. 2023. Global spread of carbapenem-resistant Enterobacteriaceae: Epidemiological features, resistance mechanisms, detection and therapy. Microbiological Research 266:127249.

7. Sabour S, Huang JY, Bhatnagar A, Gilbert SE, Karlsson M, Lonsway D, Lutgring JD, Rasheed JK, Halpin AL, Stanton RA, Gumbis S, Elkins CA, Brown AC. 2021. Detection and characterization of targeted carbapenem-resistant health care-associated threats: Findings from the Antibiotic Resistance Laboratory Network, 2017 to 2019. Antimicrobial Agents and Chemotherapy 65:10.1128/aac.01105-21.

8. CDC. Antimicrobial Resistance & Patient Safety Portal (AR&PSP) AR Lab Network Data. Atlanta, Georgia: U.S. Department of Health and Human Services, CDC. https://arpsp.cdc.gov/.

9. Hamidian M, Nigro SJ. 2019. Emergence, molecular mechanisms and global spread of carbapenem-resistant Acinetobacter baumannii. Microbial Genomics 5.

10. Afzal-Shah M, Villar HE, Livermore DM. 1999. Biochemical characteristics of a carbapenemase from an Acinetobacter baumannii isolate collected in Buenos Aires, Argentina. Journal of Antimicrobial Chemotherapy 43:127–131.

11. Afzal-Shah M, Woodford N, Livermore DM. 2001. Characterization of OXA-25, OXA-26, and OXA-27, molecular class D β-lactamases associated with carbapenem resistance in clinical isolates of Acinetobacter baumannii. Antimicrobial Agents and Chemotherapy 45:583–588.

12. Sabour S, Bantle K, Bhatnagar A, Huang JY, Biggs A, Bodnar J, Dale JL, Gleason R, Klein L, Lasure M, Lee R, Nazarian E, Schneider E, Smith L, Snippes Vagnone P, Therrien M, Tran M, Valley A, Wang C, Young EL, Lutgring JD, Brown AC. 2024. Descriptive analysis of targeted carbapenemase genes and antibiotic susceptibility profiles among carbapenem-resistant Acinetobacter baumannii tested in the Antimicrobial Resistance Laboratory Network—United States, 2017–2020. Microbiology Spectrum 12:e02828–23.

13. Tanner WD, Atkinson RM, Goel RK, Toleman MA, Benson LS, Porucznik CA, VanDerslice JA. 2017. Horizontal transfer of the bla_NDM-1_ gene to Pseudomonas aeruginosa and Acinetobacter baumannii in biofilms. FEMS Microbiology Letters 364:fnx048.

14. de Man TJB, Yaffee AQ, Zhu W, Batra D, Alyanak E, Rowe LA, McAllister G, Moulton-Meissner H, Boyd S, Flinchum A, Slayton RB, Hancock S, Spalding Walters M, Laufer Halpin A, Rasheed JK, Noble-Wang J, Kallen AJ, Limbago BM. 2021. Multispecies outbreak of Verona integron-encoded metallo-ß-lactamase-producing multidrug-resistant bacteria driven by a promiscuous incompatibility group A/C2 plasmid. Clinical Infectious Diseases 72:414–420.

15. Leski TA, Bangura U, Jimmy DH, Ansumana R, Lizewski SE, Li RW, Stenger DA, Taitt CR, Vora GJ. 2013. Identification of bla_OXA-51-like_, bla_OXA-58_, bla_DIM-1_, and bla_VIM_ carbapenemase genes in hospital Enterobacteriaceae isolates from Sierra Leone. J Clin Microbiol 51:2435–2438.

16. Manohar P, Leptihn S, Lopes BS, Nachimuthu R. 2021. Dissemination of carbapenem resistance and plasmids encoding carbapenemases in Gram-negative bacteria isolated in India. JAC-Antimicrobial Resistance 3:dlab015.

17. Long SW, Olsen RJ, Eagar TN, Beres SB, Zhao P, Davis JJ, Brettin T, Xia F, Musser JM. 2017. Population genomic analysis of 1,777 extended-spectrum beta-lactamase-producing Klebsiella pneumoniae isolates, Houston, Texas: unexpected abundance of clonal group 307. mBio 8:10.1128/mbio.00489-17.

18. Paul D, Ingti B, Bhattacharjee D, Maurya AP, Dhar D, Chakravarty A, Bhattacharjee A. 2017. An unusual occurrence of plasmid-mediated bla_OXA-23_ carbapenemase in clinical isolates of Escherichia coli from India. International Journal of Antimicrobial Agents 49:642–645.

19. Bonnet R, Marchandin H, Chanal C, Sirot D, Labia R, De Champs C, Jumas-Bilak E, Sirot J. 2002. Chromosome-encoded class D β-lactamase OXA-23 in Proteus mirabilis. Antimicrob Agents Chemother 46:2004–2006.

20. Österblad M, Karah N, Halkilahti J, Sarkkinen H, Uhlin BE, Jalava J. 2016. Rare detection of the Acinetobacter class D carbapenemase bla_OXA-23_ gene in Proteus mirabilis. Antimicrobial Agents and Chemotherapy 60:3243–3245.

21. Lange F, Pfennigwerth N, Gerigk S, Gohlke F, Oberdorfer K, Purr I, Wohanka N, Roggenkamp A, Gatermann SG, Kaase M. 2017. Dissemination of bla_OXA-58_ in Proteus mirabilis isolates from Germany. Journal of Antimicrobial Chemotherapy 72:1334–1339.

22. Leulmi Z, Kandouli C, Mihoubi I, Benlabed K, Lezzar A, Rolain J-M. 2019. First report of blaOXA-24 carbapenemase gene, armA methyltransferase and aac(6′)-Ib-cr among multidrug-resistant clinical isolates of Proteus mirabilis in Algeria. Journal of Global Antimicrobial Resistance 16:125–129.

23. Potron A, Hocquet D, Triponney P, Plésiat P, Bertrand X, Valot B. 2019. Carbapenem-susceptible OXA-23-producing Proteus mirabilis in the French community. Antimicrobial Agents and Chemotherapy 63:10.1128/aac.00191-19.

24. Bonnin RA, Girlich D, Jousset AB, Gauthier L, Cuzon G, Bogaerts P, Haenni M, Madec J-Y, Couvé-Deacon E, Barraud O, Fortineau N, Glaser P, Glupczynski Y, Dortet L, Naas T. 2020. A single Proteus mirabilis lineage from human and animal sources: a hidden reservoir of OXA-23 or OXA-58 carbapenemases in Enterobacterales. 1. Sci Rep 10:9160.

25. Hamprecht A, Sattler J, Noster J, Stelzer Y, Fuchs F, Dorth V, Gatermann SG, Göttig S. 2023. Proteus mirabilis – analysis of a concealed source of carbapenemases and development of a diagnostic algorithm for detection. Clinical Microbiology and Infection 29:1198.e1-1198.e6.

26. Shugart A, Mahon G, Huang JY, Karlsson M, Valley A, Lasure M, Gross A, Pattee B, Vaeth E, Brooks R, Maruca T, Dominguez CE, Torpey D, Francis D, Bhattarai R, Kainer MA, Chan A, Dubendris H, Greene SR, Blosser SJ, Shannon DJ, Jones K, Brennan B, Hun S, D’Angeli M, Murphy CN, Tierney M, Reese N, Bhatnagar A, Kallen A, Brown AC, Spalding Walters M. 2021. Carbapenemase production among less-common Enterobacterales genera: 10 US sites, 2018. JAC-Antimicrobial Resistance 3:dlab137.

27. Naas T, Oueslati S, Bonnin RA, Dabos ML, Zavala A, Dortet L, Retailleau P, Iorga BI. 2017. Beta-lactamase database (BLDB) – structure and function. Journal of Enzyme Inhibition and Medicinal Chemistry 32:917–919.

28. Krawczyk PS, Lipinski L, Dziembowski A. 2018. PlasFlow: predicting plasmid sequences in metagenomic data using genome signatures. Nucleic Acids Research 46:e35.

29. Wick RR, Judd LM, Gorrie CL, Holt KE. 2017. Unicycler: Resolving bacterial genome assemblies from short and long sequencing reads. PLOS Computational Biology 13:e1005595.

30. Carattoli A, Hasman H. 2020. PlasmidFinder and In Silico pMLST: Identification and typing of plasmid replicons in whole-genome sequencing (WGS), p. 285–294. In de la Cruz, F (ed.), Horizontal Gene Transfer: Methods and Protocols. Springer US, New York, NY.

31. Altschul SF, Gish W, Miller W, Myers EW, Lipman DJ. 1990. Basic local alignment search tool. Journal of Molecular Biology 215:403–410.

32. Sayers EW, Beck J, Brister JR, Bolton EE, Canese K, Comeau DC, Funk K, Ketter A, Kim S, Kimchi A, Kitts PA, Kuznetsov A, Lathrop S, Lu Z, McGarvey K, Madden TL, Murphy TD, O’Leary N, Phan L, Schneider VA, Thibaud-Nissen F, Trawick BW, Pruitt KD, Ostell J. 2020. Database resources of the National Center for Biotechnology Information. Nucleic Acids Research 48:D9–D16.

33. Alikhan N-F, Petty NK, Ben Zakour NL, Beatson SA. 2011. BLAST ring image generator (BRIG): simple prokaryote genome comparisons. BMC Genomics 12:402.

34. Lutgring JD, Balbuena R, Reese N, Gilbert SE, Ansari U, Bhatnagar A, Boyd S, Campbell D, Cochran J, Haynie J, Ilutsik J, Longo C, Swint S, Rasheed JK, Brown AC, Karlsson M. 2020. Antibiotic susceptibility of NDM-producing Enterobacterales collected in the United States in 2017 and 2018. Antimicrobial Agents and Chemotherapy 64:10.1128/aac.00499-20.

35. CLSI. 2022. Performance standards for antimicrobial susceptibility testing. 32 ed. CLSI supplement M100. Clinical and Laboratory Standards Institute.

36. U.S. Food and Drug Administration. Antibacterial susceptibility test interpretive criteria. https://www.fda.gov/drugs/development-resources/antibacterial-susceptibility-test-interpretive-criteria.

37. Poirel L, Potron A, Nordmann P. 2012. OXA-48-like carbapenemases: the phantom menace. Journal of Antimicrobial Chemotherapy 67:1597–1606.

38. Fursova NK, Astashkin EI, Knyazeva AI, Kartsev NN, Leonova ES, Ershova ON, Alexandrova IA, Kurdyumova NV, Sazikina SYu, Volozhantsev NV, Svetoch EA, Dyatlov IA. 2015. The spread of blaOXA-48 and blaOXA-244 carbapenemase genes among Klebsiella pneumoniae, Proteus mirabilis and Enterobacter spp. isolated in Moscow, Russia. Annals of Clinical Microbiology and Antimicrobials 14:46.

39. Pitout JDD, Peirano G, Kock MM, Strydom K-A, Matsumura Y. 2019. The global ascendency of OXA-48-type carbapenemases. Clinical Microbiology Reviews 33:10.1128/cmr.00102-19.

40. Pedraza R, Kieffer N, Guzmán-Puche J, Artacho MJ, Pitart C, Hernández-García M, Vila J, Cantón R, Martinez-Martinez L. 2022. Hidden dissemination of carbapenem-susceptible OXA-48-producing Proteus mirabilis. Journal of Antimicrobial Chemotherapy 77:3009–3015.

41. Boyd SE, Holmes A, Peck R, Livermore DM, Hope W. 2022. OXA-48-like β-lactamases: global epidemiology, treatment options, and development pipeline. Antimicrobial Agents and Chemotherapy 66:e00216–22.

42. Conlan S, Park M, Deming C, Thomas PJ, Young AC, Coleman H, Sison C, Weingarten RA, Lau AF, Dekker JP, Palmore TN, Frank KM, Segre JA. 2016. Plasmid dynamics in KPC-positive Klebsiella pneumoniae during long-term patient colonization. mBio 7:e00742–16.

43. Shahid M, Ahmad N, Saeed NK, Shadab M, Joji RM, Al-Mahmeed A, Bindayna KM, Tabbara KS, Dar FK. 2022. Clinical carbapenem-resistant Klebsiella pneumoniae isolates simultaneously harboring bla_NDM-1_, bla_OXA_ types and qnrS genes from the Kingdom of Bahrain: Resistance profile and genetic environment. Frontiers in Cellular and Infection Microbiology 12.

44. La M-V, Jureen R, Lin RTP, Teo JWP. 2014. Unusual detection of an Acinetobacter Class D carbapenemase gene, bla_OXA-23_, in a clinical Escherichia coli isolate. J Clin Microbiol 52:3822–3823.

45. Nigro SJ, Hall RM. 2016. Structure and context of Acinetobacter transposons carrying the oxa23 carbapenemase gene. Journal of Antimicrobial Chemotherapy 71:1135–1147.

46. Octavia S, Xu W, Ng OT, Marimuthu K, Venkatachalam I, Cheng B, Lin RTP, Teo JWP. 2020. Identification of AbaR4 Acinetobacter baumannii resistance island in clinical isolates of bla_OXA-23_-positive Proteus mirabilis. Journal of Antimicrobial Chemotherapy 75:521–525.

47. Bi D, Xie R, Zheng J, Yang H, Zhu X, Ou H-Y, Wei Q. 2019. Large-scale identification of AbaR-type genomic islands in Acinetobacter baumannii reveals diverse insertion sites and clonal lineage-specific antimicrobial resistance gene profiles. Antimicrobial Agents and Chemotherapy 63:10.1128/aac.02526-18.

48. Marquez-Ortiz RA, Haggerty L, Olarte N, Duarte C, Garza-Ramos U, Silva-Sanchez J, Castro BE, Sim EM, Beltran M, Moncada MV, Valderrama A, Castellanos JE, Charles IG, Vanegas N, Escobar-Perez J, Petty NK. 2017. Genomic epidemiology of NDM-1-encoding plasmids in Latin American clinical isolates reveals insights into the evolution of multidrug resistance. Genome Biology and Evolution 9:1725–1741.

49. Chen Z, Li H, Feng J, Li Y, Chen X, Guo X, Chen W, Wang L, Lin L, Yang H, Yang W, Wang J, Zhou D, Liu C, Yin Z. 2015. NDM-1 encoded by a pNDM-BJ01-like plasmid p3SP-NDM in clinical Enterobacter aerogenes. Frontiers in Microbiology 6.

50. Ambrose SJ, Hamidian M, Hall RM. 2022. Extensively resistant Acinetobacter baumannii isolate RCH52 carries several resistance genes derived from an IncC plasmid. Journal of Antimicrobial Chemotherapy 77:930–933.

