## Supplemental Figure 1 for "Evaluation of Enterobacterales carrying *Acinetobacter*-associated *bla*_OXA_ genes—United States, 2017–2022"

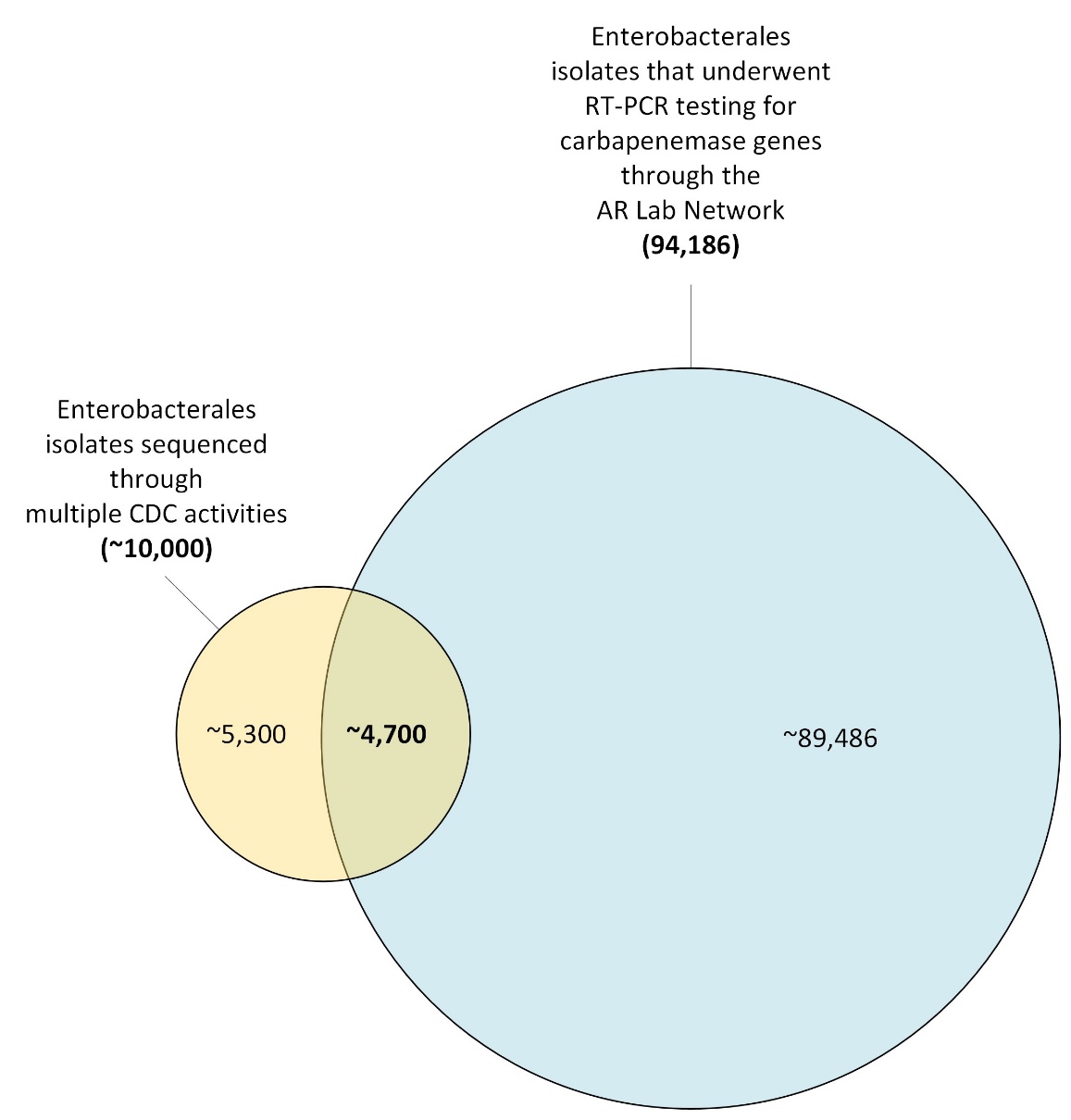

**Figure S1. Illustration of isolate data analyzed in this evaluation.** The Venn diagram represents the denominators included in the analyses. Key denominators referenced in the main text are shown in bold. As part of AR Lab Network testing from 2017–2022, 94,186 Enterobacterales isolates underwent RT-PCR testing for one of the five targeted carbapenemase genes (*bla*_KPC_, *bla*_NDM_, *bla*_OXA-48-like_, *bla*_IMP_, and *bla*_VIM_), represented by the large, blue circle. Of those 94,186 isolates, ~4,700 isolates also underwent whole-genome sequencing (the intersection of the two circles). The small, yellow circle represents the ~10,000 Enterobacterales isolates that were collected and sequenced through multiple CDC activities (surveillance activities, outbreak investigations, reference testing, and AR Lab Network activities) from 2017–2022. The current state of our data infrastructure and challenges in linking data from different sources makes it infeasible to determine what proportion of the ~10,000 isolate sequences are duplicates due to resequencing, and thus provide precise denominators for isolates sequenced through each activity. Abbreviations: AR Lab Network, Antimicrobial Resistance Laboratory Network; RT-PCR, real-time-PCR; CDC, Centers for Disease Control and Prevention.
